# Serial dependence tracks objects and scenes in parallel and independently

**DOI:** 10.1101/2021.07.12.452024

**Authors:** Thérèse Collins

## Abstract

The visual world is made up of objects and scenes. Object perception requires both discriminating an individual object from others and binding together different perceptual samples of that object across time. Such binding manifests by serial dependence, the attraction of the current perception of a visual attribute towards values of that attribute seen in the recent past. Scene perception is subserved by global mechanisms like ensemble perception, the rapid extraction of the average feature value of a group of objects. The current study examined to what extent the perception of single objects in multi-object scenes depended on previous feature values of that object, or on the average previous attribute of all objects in the scene. Results show that serial dependence occurs independently on two simultaneously present objects, that ensemble perception depends only on previous ensembles, and that serial dependence of an individual object occurs only on the features of that particular object. These results suggest that the temporal integration of successive perceptual samples operates simultaneously at independent levels of visual processing.

## Introduction

The visual world as we perceive it is made up of objects and scenes. Perceiving individual objects requires both discriminating them from others and properly binding together different perceptual samples of the same object across viewpoints, retinal locations, changes in ambient lighting, and time. Temporal binding is thought to be subserved by a mechanism called a continuity field, a spatio-temporally tuned operator that integrates successive perceptual samples over a brief moment in time and across a restricted region of space. Continuity fields may promote perceptual stability across time by smoothing out spurious fluctuations in the proximal stimulus. Continuity fields manifest in the laboratory as a phenomenon called serial dependence, small misperceptions of currently presented visual attributes as pulled towards values of that attribute seen in the immediate past. Perceiving scenes is aided by mechanisms that extract summary statistics and average them into global representations of cluttered environments ((Alvarez, 2011)). Observers are remarkably good at extracting the average value of a group of objects, such as the average orientation of a group of Gabor patches, the average speed and direction of moving dots, or the average emotional expression in a crowd of faces (e.g. (Haberman & Whitney, 2007)). Importantly, the attributes of the individual objects that make up a group are often less well perceived than the average attribute ((Ariely, 2001)).

The two effects may be contradictory: the attraction of object perception towards features values in its recent past versus the perceptual predominance of the average attribute. It remains unknown how individual object and ensemble representations are integrated across time.

Serial dependence has been examined mainly in the context of sequential presentations of a single object, and observer reports of a feature of that object. It is thus unknown to what extent the perception of individual objects in multi-object scenes depends on previous ensemble representations of the scene, or whether independent representations can be maintained for individual objects and global scene characteristics. Ensemble perception is subject to serial dependence: when asked to report the average attribute of a set of stimuli, subject responses were pulled towards the average of the previous set ((Manassi et al., 2017)). Manassi et al. (2017) also showed that serial dependence occurred between single objects and ensembles, and vice versa, but they did not contrast the effect of previous individual objects versus previous ensembles. This is the crucial question to determine how object and scene representations interact to ensure perceptual stability across time.

The main question of interest here is whether an individual object that has been averaged into an ensemble representation can still influence upcoming perception of the individual item. If that were the case, it would suggest that multiple continuity fields are maintained simultaneously for objects and for the ensemble representation including those objects. This seems unlikely, given the literature on how representations of individual objects in visual short-term memory is biased by the summary statistics of the current scene ((Brady & Alvarez, 2011; Lew & Vul, 2015)). Furthermore, long-term memories may resemble ensemble representations ((Richards et al., 2014)) and scene information is a powerful cue for long-term object recall (e.g. (Brady et al., 2011; Hollingworth, 2006)).

The current study examined to what extent the current perception of a single shape and the current perception of the average of a set of shapes depended on previously seen shapes or ensembles of shapes (Figure 1). The previous literature on ensemble perception and its influence on memory for individual objects suggests the following hypothesis: that both the current perception of an individual shape within a scene and the current perception of the average shape will be pulled towards the average shape of previous scenes.

**Figure 1.**
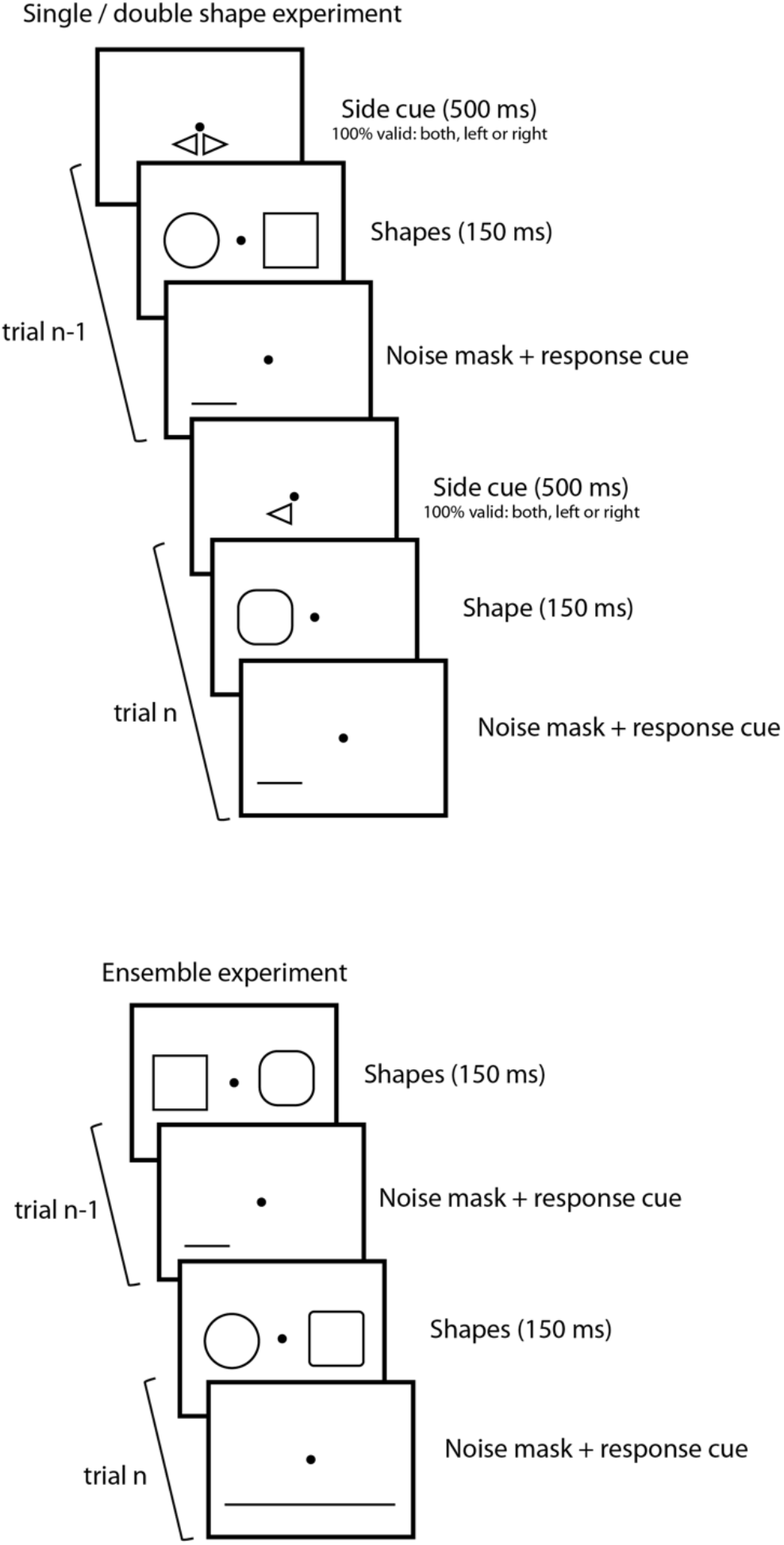
Procedure. **Top panel:** single / double shape experiment (Exp. 1). Two trials are pictured. In each, a 500-ms cue indicates number of shapes (one or two) and, in the case of one shape, its side (left or right). Shapes are then presented for 150 ms, followed by a noise mask and response cue until subject response. **Bottom panel:** ensemble experiment (Exp. 2). There were always two shapes, presented for 150 ms, followed by the noise mask and response cue until subject response.

## Materials and methods

### Participants

Fifteen subjects participated in Experiment 1 (2 women, mean age 29 years old, range 18-65). Nineteen subjects participated in Experiment 2 (3 women, mean age 33 years old, range 18-56). The experiments were run on the online platform testable.org, and subjects were recruited through the Testable Minds subject pool. The number of subjects was determined by a power analysis based on reports of serial dependence effect sizes in the literature, which places the number at around 5 subjects for a power of 99% (i.e. estimated 1% probability of a false negative). The number of subjects was increased because it was practically feasible and ensured greater power, given that some of the conditions tested here were expected to have smaller serial dependence than what has been reported before. The relevant ethical information pertaining to the laboratory version of the study and approved by the French national ethics committee (CPP) in accordance with the declaration of Helsinki was provided at the onset of the experiment, and participants had to click to accept the consent form before proceeding with the experiment. Sex, age and screen characteristics, but no personally identifying information, were recorded for each participant.

### Stimuli

Because the experiment was run on an online platform, exact control of stimulus size was not possible. Testable.org displays stimuli as images centered on observers’ personal computer screens, adjusted for each participant’s screen resolution but without distorting the image proportions (i.e. the same adjustment in width and height). Observers were instructed to sit 60 cm (arm’s length) from their screen. All stimuli were 1024×768 pixel images; stimulus sizes below are given for a typical laptop screen, but because of individual screens the values can vary from 0.7 to 1.2 in proportion. Stimuli were generated using Psychtoolbox for Matlab ((Brainard, D., 1997; Kleiner, M et al., 2007; Pelli, G., 1997)) and saved as images.

A black dot, 0.25 degrees of visual angle (dva) in diameter and presented at screen center, served as a fixation point. Test stimuli were 100 geometrical shapes that ranged from a circle to a square. For trials with two shapes, there were thus 10^4^ possible combinations; a random subset of 1000 trials was selected for Experiment 1 and another random subset of 1800 trials for Experiment 2. Shapes were 2.7 dva wide (circle diameter or square side). The transition from one shape to the other was made by placing, at each corner of a square, arcs defined by a central angle from 45° (circle) to 0° (square), in steps of 0.45°. For ease of interpretation, each shape was given a value from 0 (circle) to 1 (square); the fully ambiguous shape (value of 0.50) was not included because there is no correct response. Shapes were embedded in pixel noise. Masks were screens of pixel noise alone.

### Procedure

Experiment 1 had two parts. The first 400 trials of the experiment were single shape trials. On each trial, a shape appeared at screen center for 150 ms, followed by a mask which also served as the response screen. The second part of the experiment was the double shape session, composed of 1000 double shape trials and 400 single shape trials (200 shapes on the left, 200 shapes on the right), randomly interleaved. The subject was cued about the number of shapes and, if there was only one, about its location (left or right). The cue was to ensure that any differences between the single-shape condition in blocked versus interleaved conditions could not be attributed to uncertainty about target location or reduced attention (the strength of serial dependence decreases in the absence of attention; (Fischer & Whitney, 2014)). In double-shape trials, two shapes appeared to the left and right of fixation for 150 ms, followed by the mask. A response cue appeared simultaneously with the mask and indicated the shape the subject was to report (left or right). Half of the trials called for a response about the shape on the left, half about the shape on the right. Subjects responded by pressing one of two buttons on the keyboard (one for “circle” and another for “square”), followed immediately by the cue for the next trial. In single shape trials, the procedure was identical except that only one shape appeared, on the left or right, and the response cue was always valid.

Experiment 2 was similar except that in addition to reporting one of the two shapes (identically to the double-shape trials in Experiment 1), on most trials participants had to report the average of the two shapes. In this case, the response cue underlined both shapes. There was no pre-cue since there were always two shapes on screen. There were 516 single-report trials (29%) and 1284 ensemble-report trials, randomly interleaved.

### Data analysis

To quantify performance, psychometric functions were fit on individual response data using the quickpsy package developed for R by ((Linares & López-Moliner, 2016)). Two parameters are of interest: the slope of the psychometric function, and the point of subjective equality (PSE), i.e. the stimulus level for which subjects selected each response option 50% of the time. Average psychometric functions shown in figures were obtained by bootstrapping individual fits.

Serial dependence was calculated as a percentage by considering error trials (i.e. trials in which the shape value was less than 0.5 in which observers reported seeing a square, and trials in which the shape value was greater than 0.5 in which observers reported seeing a circle). The number of trials in which the response erred towards the previously presented shape was divided by the total number of errors. If there was no relationship between current response and previous shape, 50% of the errors should be preceded by more circle-like shapes, and 50% by more square-like shapes. For ease of interpretation, 50% was thus subtracted from the percent serial dependence. Thus, no serial dependence is 0%, 50% serial dependence is if all erroneous responses erred towards the previous shape, and −50% serial dependence is if all erroneous responses erred away from the previous shape (a repulsive effect).

Temporal tuning was described by fitting an exponential function of the form a*exp(-b*x) to the percent serial dependence on trials further and further into the past (from 1 trial back to 10 trials back). The significance of parameters a and b is an indicator of whether an exponential is a good description of the pattern.

## Results

Subjects were good at reporting single shapes and there was no cost of interleaving single and double shape trials. Figure 2 shows psychometric functions for the two single-shape conditions (blocked and interleaved with double-shape trials). There were no significant differences in performance between them, on either slope or point of subjective equality (PSE) as shown by the overlapping 95% confidence intervals (insets in Figure 2).

**Figure 2.**
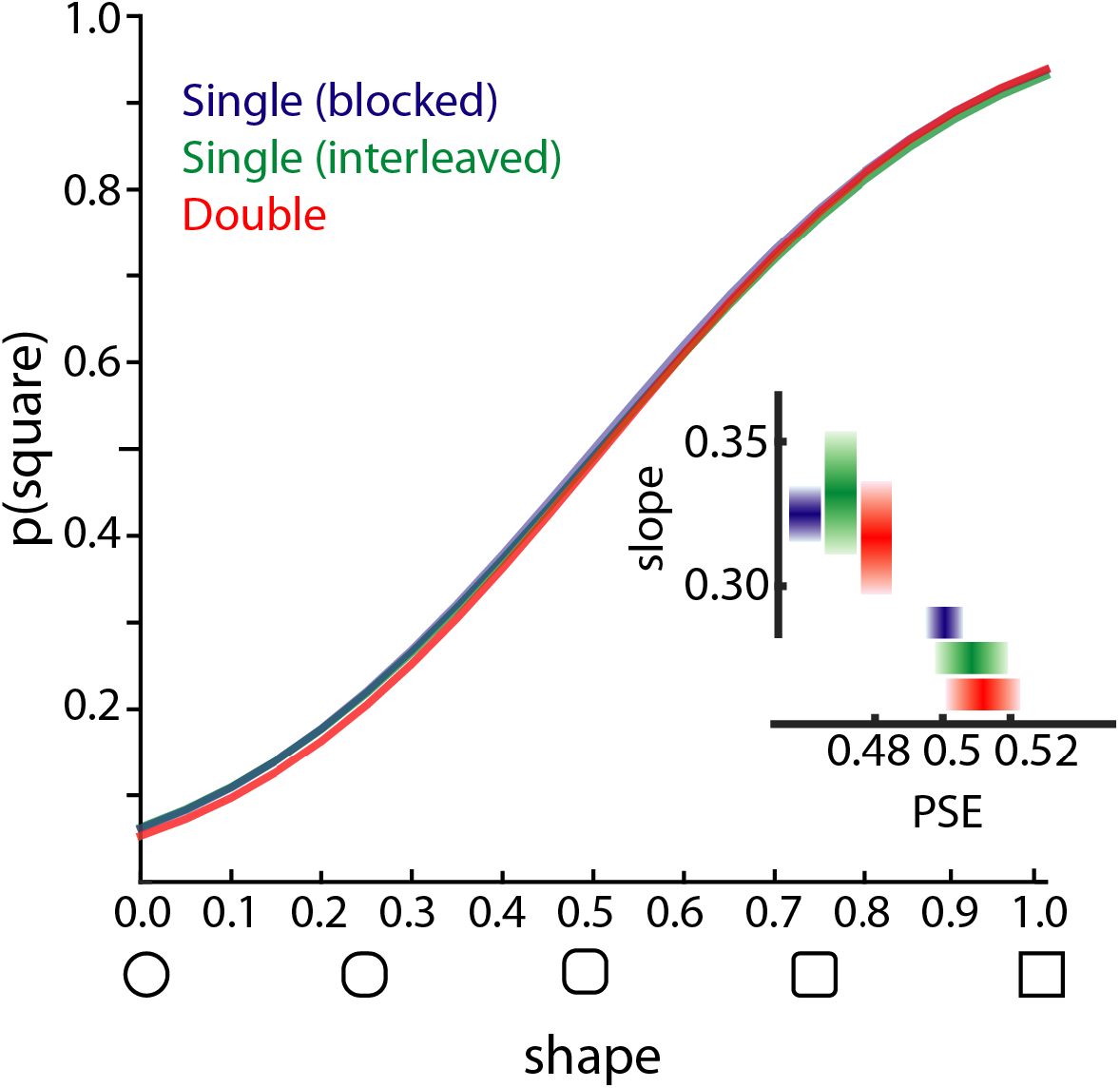
Psychometric functions (probability of responding “square” as a function of physical shape value), in the single/double shape experiment (Exp. 1). Blue: single shape trials, blocked design. Green: single shape trials, interleaved with double shape trials. Red: double-shape trials. Inset: Average slope and PSE ( ± bootstrapped 95% confidence intervals).

Figure 3 quantifies serial dependence for different conditions. The shaded bar surrounding the average corresponds to 95% confidence intervals, based on bootstrapping individual data, such that any condition for which the lower bound of the confidence interval is above zero is considered significant. Perception of single shapes depended on the immediately preceding shape, in both blocked and interleaved conditions (blocked: 6.1%, 95% CI: [3.8-8.5%]; interleaved: 4.0%, [0.6-7.3%]). The rightmost bars in Figure 3 show the (non-significant) serial dependence of current response on future shape. This analysis is a control, since of course there can be no influence of the future on the present.

**Figure 3.**
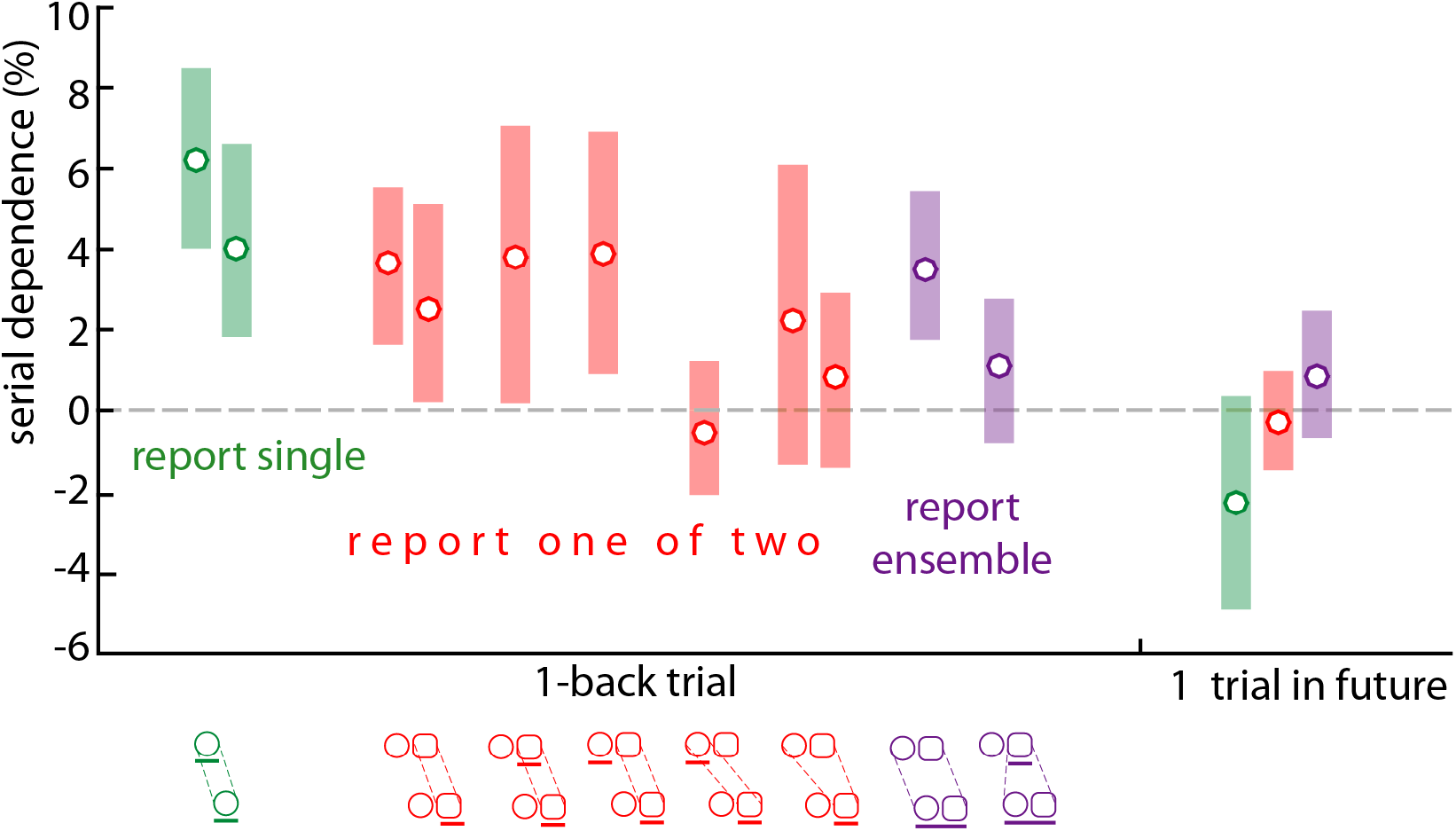
Percent serial dependence on the 1-back and +1 trials as a function of report (one, two or average), in Experiments 1 and 2. The conditions, in order from left to right, are as follows. Green: single shapes in blocked and interleaved designs. Red: double shape trials in which subjects reported only one of the two shapes, serial dependence on the previous shape on the same side for all double-shape trials (data from Exp. 1 & 2); SD on previous shape on the same side when it was cued (Exp. 1); SD on previous shape on the same side when it was uncued (Exp. 1); SD on previous cued shape on the opposite side (Exp. 1); SD on the average of the two previous shapes (Exp. 1 & 2). Purple (Exp. 2): double-shape trials in which subjects reported the average of the two shapes: SD on the average of the two previous shapes; SD on the previously cued single shape.

Perception of single shapes when two shapes were present was comparable to perception of single shapes presented alone. The psychometric function is shown in Figure 2 and overlaps nicely with the two single-shape conditions. Perception of single shape in a double-shape trial depended on the immediately preceding shape (3.6%, 95% CI: [1.6– 5.4%]), and this condition was replicated in Exp. 2 (2.7%, [0.4%-4.9%]). In the double-shape trials, it is possible to look at the influence of the preceding shape on the same side depending on whether that shape was cued or not. Figure 3 breaks down serial dependence in the double-shape trials as a function of whether the previous shape at the same location was cued or not. Current response depended on previous shape at the same location, but it did not matter very much if that shape had been cued or not (cued: 3.3%, [0.5–6.1%]; uncued: 3.8%, [0.7–7.1%]). However, the serial dependence of current response on previously cued shape was not significantly different from zero when the previously cued shape was at the opposite location (0.6%, [−1.6–2.6%]). These results suggest that each object (or side) is monitored independently.

Figure 3 also illustrates how current perception depends on the average shape seen in the previous trial. This analysis considers both single-shape and double-shape trials that were preceded by a double-shape trial. The amount of serial dependence was not significantly different from zero (2.2%, [−0.2–6.0%]) and this condition was replicated in Exp. 2 (0.8%, [−1.4–3.0%)). This suggests that ensemble perception is not obligatory: when subjects reported individual items, their responses were independent from the previous average visual attribute.

Serial dependence is temporally tuned, with maximal dependence of current perception on the immediately preceding trial and gradually decreasing dependence on trials farther into the past. Such a pattern was replicated here (Figure 4) for all conditions. Exponentials with significant a and b parameters are in thick lines in Figure 4; thin lines denote non-significant parameters.

**Figure 4.**
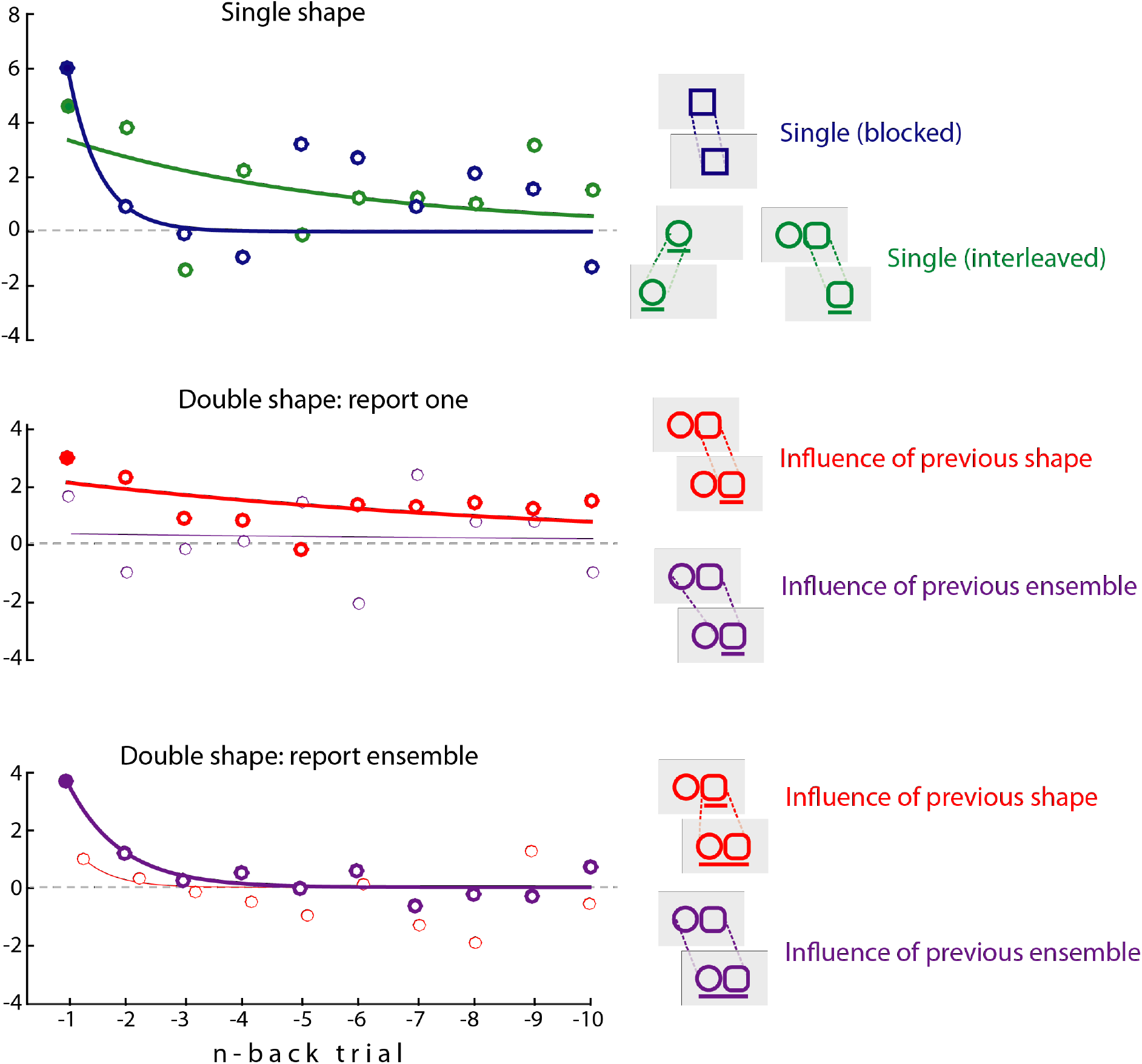
Percent serial dependence. Data for the 1-back trial is identical to that reported in Figure 3. Filled circles indicate significance. Thick lines represent significant exponential fits, thin line non-significant. **Top panel** (Experiment 1): Single shape trials, blocked (blue) or interleaved (green) design. **Middle panel** (Experiment 1): Double-shape trials in which subjects had to report one of the two shapes. In red, influence of the previous shape on the same side. In purple, influence of the previous ensemble. **Bottom panel** (Experiment 2): Double-shape trials in which subjects had to report the average of the two shapes. In red, influence of the shape on the previously cued side. In purple, influence of the previous ensemble.

To further explore serial dependence of individual versus ensemble features, Experiment 2 required a new group of participants (n=19) to report the average of the two shapes. Stimuli were identical to the first experiment except for the response cue which underlined both shapes when an average response was required (Figure 1). For comparison purposes, these ensemble-shape trials were interleaved with double-shape trials, i.e. trials in which two shapes were present but subjects reported the shape of either the left or right shape, as instructed by a response cue, just like in the first experiment.

Figure 5 shows psychometric functions for single and ensemble responses in the second experiment, with bootstrapped mean and 95% confidence intervals in the inset. Overlapping confidence intervals show that there were no significant differences between the two conditions: subjects were as good at reporting one of two shapes as the average of two shapes. This confirms many previous reports of ensemble perception in the literature.

**Figure 5.**
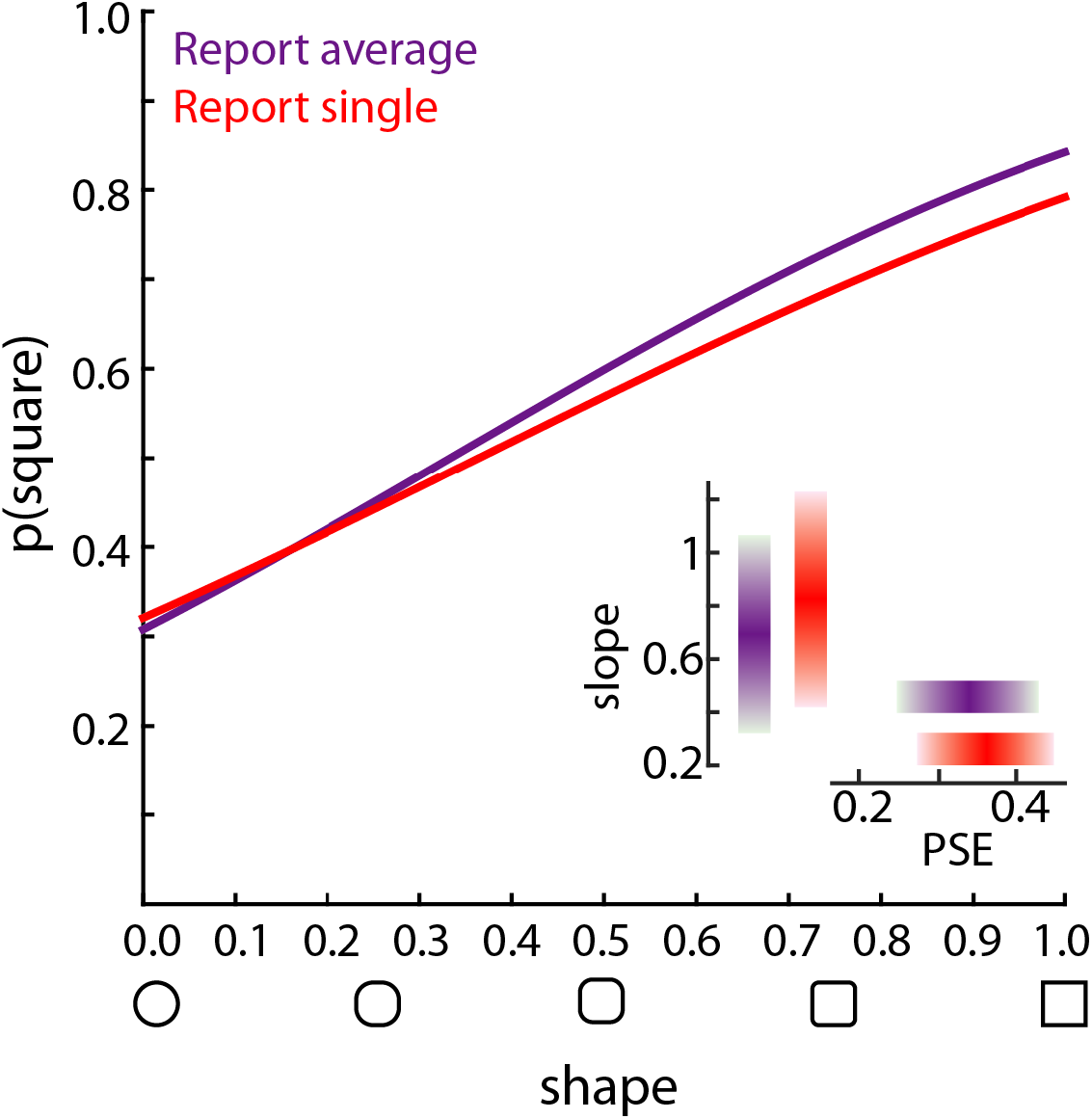
Psychometric functions (probability of responding “square” as a function of physical shape value), in the ensemble experiment (Exp. 2). Purple: double-shape trials in which subjects reported the average shape. Red: double-shape trials in which subjects reported one of the two shapes. Inset: Average slope and PSE ( ± bootstrapped 95% confidence intervals).

Percent serial dependence was calculated as before (Figure 3). When participants had to report the average of two shapes, their perception depended on the previous ensemble (3.1%, [1.6%−4.7%]). The temporal profile was well described by an exponential (Figure 4). However, their perception did not depend significantly on the shape on the previously cued side (1.1%, [−0.7%−2.8%]).

## Discussion

When participants were asked to report the shape of an individual object, their responses were attracted towards the shape of that object seen in the recent past. They were not influenced by a shape on the opposite side, even when that shape had been previously cued for report. This serial dependence between objects at the same location, but not across locations, may seem to contrast with results from other studies showing that serial dependence operates across locations (if they are not too far apart;(Collins, 2019)). However, in these previous studies, there was only one object, compared with the two in the current experiment. This result suggests that individual objects within multi-object scenes are monitored by independent continuity fields.

Contrary to the initial hypothesis, the perception of individual object shape was not significantly influenced by the average shape of the previous multi-shape scene, suggesting that ensemble perception does not obligatorily influence upcoming individual shape perception.

Finally, when participants were asked to report the average of two shapes, they were influenced by the previous ensemble representation (replicating Manassi et al., 2017). There was no influence of previously reported individual shapes on ensemble responses, in direct contrast to results reported by Manassi et al. (2017). The key difference may be that in the current study, there were always multiple objects, whereas in Manassi et al (2017), ensemble perception was influenced by single shapes (presented alone). Furthermore, Manassi et al. (2017) did not ask participants to report one object in a multi-object scene, and they did not examine to what extent single-object responses were attracted to the previous average or to the previous feature at the same location.

Taken together, the results of the current show that the object representations that drive perception are influenced by multiple aspects of stimulus history, depending on the goal of current perception: individual object or global scene perception. The perception of individual objects in multi-object scenes is attracted towards previous features of that object, but not towards the ensemble representation of the previous scene, or by the previous features of a nearby object. Ensemble perception is influenced by previous ensembles, but not by the features of individual objects in a previously viewed scene. Because observers were not told whether they would have to report one of the objects or the average until after having viewed the scene, this means that single object representations and ensemble representations are maintained and integrated with previous history in parallel. In other words, multiple continuity fields operate simultaneously on object and scene representations.

